# High fidelity lineage tracing in mouse pre-implantation embryos using primed conversion of photoconvertible proteins

**DOI:** 10.1101/218354

**Authors:** Maaike Welling, Manuel Alexander Mohr, Aaron Ponti, Lluc Rullan Sabater, Andrea Boni, Prisca Liberali, Pawel Pelczar, Periklis Pantazis

## Abstract

Accurate lineage reconstruction of mammalian pre-implantation development is essential for inferring the earliest cell fate decisions of mammalian development. Lineage tracing using global labeling techniques is complicated by increasing cell density and rapid embryo rotation, impeding automatic alignment and rendering accurate cell tracking of obtained four-dimensional imaging data sets highly challenging. Here, we exploit the advantageous properties of primed convertible fluorescent proteins (pr-pcFPs) to simultaneously visualize the global green and the photoconverted red population to minimize tracking uncertainties over prolonged time windows. Confined primed conversion of H2B-pr-mEosFP labeled nuclei combined with light-sheet imaging greatly facilitates segmentation, classification, and tracking of individual nuclei from the 4-cell stage up to the blastocyst. Using green and red labels as fiducial markers, we computationally correct for rotational and translational drift and accomplish high fidelity lineage tracing combined with a reduced data size – addressing majors concerns in the field of volumetric embryo imaging.

## Introduction

Accurate lineage tracing and precise tracking of single cells in pre-implantation embryos is essential for a mechanistic understanding of the first cell fate decisions during mammalian development^1^. Selective plane illumination microscopy (SPIM) has the potential to play a major role in achieving comprehensive, non-invasive imaging of mammalian pre-implantation development. During these early steps of development, a major fraction of embryos (n=5/11, 45% in this study) exhibit confounding rotational and translational drift (Videos 1 and 2), which often leads researchers to exclude these embryos from analysis, drastically decreasing efficiency, losing valuable data, and potentially biasing downstream results^2,3^. While high imaging rates have helped to overcome these challenges for samples like zebrafish embryos, they demand increased data storage capacities. Higher framerates can further increase photodamage from laser overexposure and are hence less applicable for highly sensitive mouse embryos^2,4^.

Sparse labeling strategies using green-to-red photoconvertible fluorescent proteins (pcFPs) merit a great potential for facilitating lineage tracing and trophectoderm (TE) and inner-cell-mass (ICM) fate assignments after photoconversion^5^. However, to our knowledge these sparse labels have not been combined with SPIM - presumably because photoconversion has been limited by the need for axially unconfined, potentially photodamaging, intense violet light^6^. Our recent report of a novel photochemical mechanism called “primed conversion” overcomes this long-standing problem by using dual-wavelength illumination with blue 488nm, and far-red 730nm laser light instead. Importantly, primed conversion allows for confined conversion of small volumes in three dimensions (3D) by selectively intersecting the two laser beams in a common focal spot, yielding axial confinement unachievable using 405nm photoconversion^7,8^. While primed conversion was previously only reported for Dendra2, the discovery of the mechanism responsible for primed conversion enabled the rational engineering of primed convertible (“pr-”) variants of most pcFPs^9^. Consequently, we found that pr-pcFP variants based on the Eos-family of *Anthozoa*-derived pcFPs undergo primed conversion efficiently and exhibit high levels of brightness and photostability, essential properties for long-term imaging in a SPIM^9^.

Here we show that primed conversion of single cells in early stages of mouse development allows for computational correction of spatial and rotational drift, which minimizes tracking and lineage tracing uncertainties over prolonged time windows.

## Results and Discussion

### H2B-pr-mEosFP labeled cells primed converted at the 4-cell stage can be visualized up to the blastocyst stage

In order to assess which protein of the Eos-family is most suitable for long-term cell tracking and lineage tracing experiments in mouse embryos, we directly compared pr-mEos2 and pr-mEosFP. We injected mouse zygotes with mRNAs encoding for the histone fusions H2B-pr-mEos2 or H2B-pr-mEosFP and imaged them at different developmental stages to observe potential detrimental effects. Embryos injected with mRNA encoding for H2B-pr-mEosFP showed no visible signs of developmental impairment, similar to un-injected control embryos (Figure 1 – figure supplement 1a). In contrast, H2B-pr-mEos2 injected embryos showed partly divided, seemingly connected nuclei and prematurely arrested in development (n=30/30) (Figure 1 – figure supplement 1b). This apparent inability to separate the nuclei during cell division is likely due to a residual tendency of mEos2 to oligomerize, as proposed previously^10^ (Figure 1 – figure supplement 1). As a consequence, we identified primed convertible mEosFP (pr-mEosFP) as the optimal fluorescent protein variant for *in vivo* primed conversion followed by long-term imaging.

**Figure 1.**
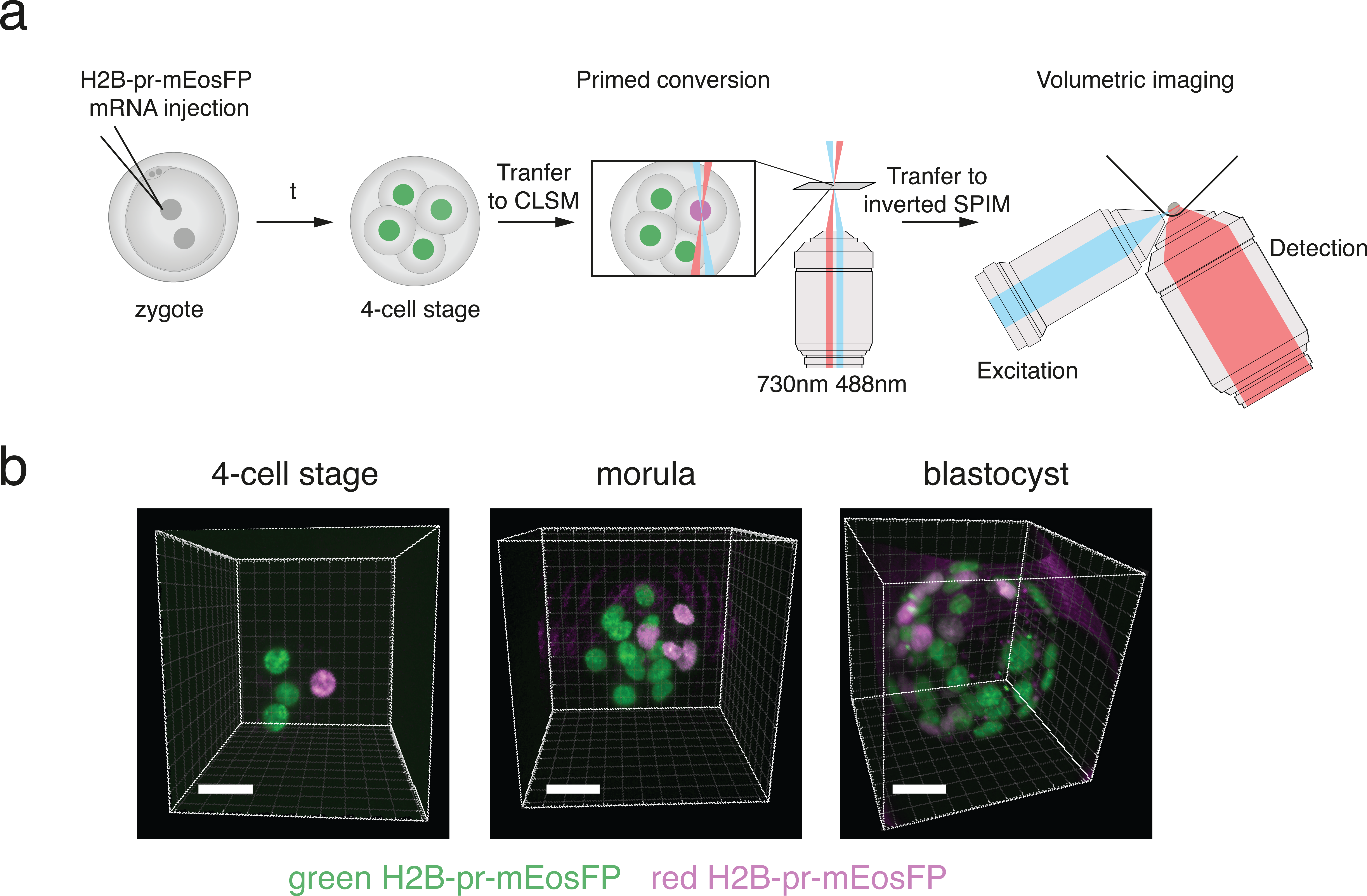
H2B-pr-mEosFP injected embryos develop to the blastocyst stage. (**a**) Experimental setup: Zygotes are injected with H2B-pr-mEosFP mRNA. At the 4-cell stage confined primed conversion of a single nucleus is performed using intersecting 488nm and 730nm lasers. The embryos are transferred to an inverted SPIM for non-invasive imaging of their development up to the blastocyst stage. Images are taken every 7.5 or 15 minutes. (**b**) Embryos injected with mRNA encoding H2B-pr-mEosFP and converted at the 4-cell-stage develop normally and maintain visibility of the red label up to the early blastocyst stage. pr-mEosFP fluorescence (green) and primed converted pr-mEosFP fluorescence (magenta). N ≥ 200 out of ≥ 10 independent experiments. Scale bar, 20 µm.

Next, we investigated whether a single round of green-to-red photoconversion at the 4-cell stage would create a sufficiently large pool of red-converted protein that could be followed throughout development until the early blastocyst stage. For this purpose, we used confined primed conversion to photoconvert a single nucleus of an H2B-pr-mEosFP expressing embryo at the 4-cell stage^8^, and monitored early embryo development for 60 hours in an inverted SPIM^2^ (Figure 1a). Photoconverted embryos developed healthily and the red daughter cells of the initially primed converted single cell were clearly distinguishable from their non-converted green counterparts up to the blastocyst stage (Figure 1b; Figure 1 – supplement 2).

### Dual labeling of pre-implantation embryos greatly facilitates automatic segmentation, tracking and lineage tracing

The observation that cells converted at the 4-cell stage can be visualized up to the blastocyst stage prompted us to ask if sparsely labeled subsets of cells could aid computational reorientation and automated lineage tracing in embryos that exhibit dramatic rotational and translational drift, (Videos 1 and 2). To this end, we developed a computational pipeline for automated segmentation, cell tracking, and lineage tracing. This algorithm uniquely takes advantage of the sparse red population to correct for translational and rotational drift as well as to simplify lineage reconstruction (Figure 2a). In the 5-dimensional (5D, i.e. 3 spatial dimensions, time, color) imaging data, cells were first segmented based on size, shape, and fluorescence taking into account both color channels. Specifically, as the red signal diminishes over time while red background autofluorescence increases, the dual labeling enables the identification of weaker fluorescent red nuclei at late time points by their overlap with green signal that has lower autofluorescence (Figure 2a, left column). Also, the dual color information allows for cell distinction in instances otherwise rendered ambiguous through high cell density and proximity of nuclei. For instance, we were able to distinguish nuclei that would have been identified as a single nucleus even after manual validation (Figure 2 – figure supplement 1a-c).

**Figure 2.**
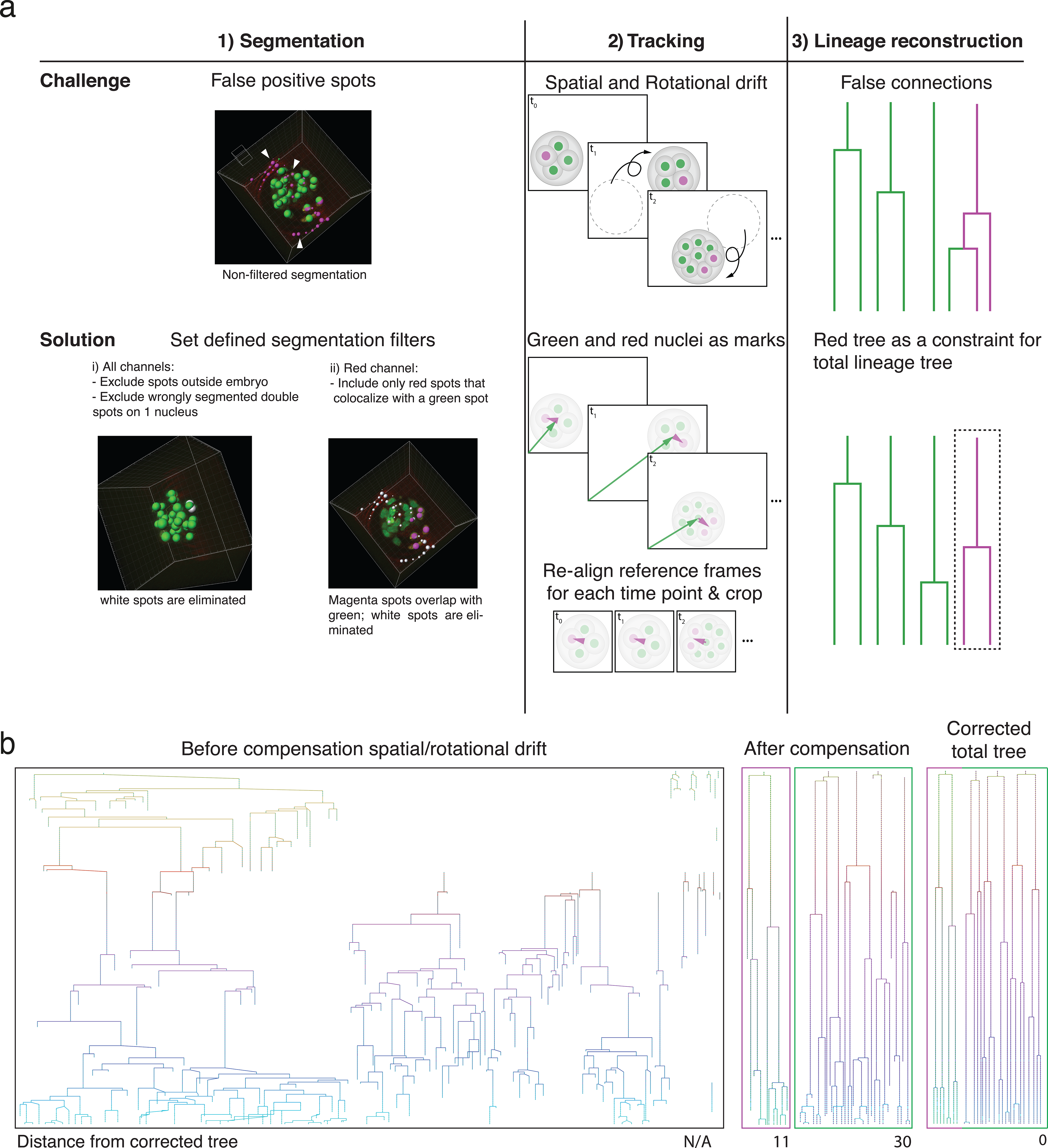
An automated segmentation, tracking, and lineage tracing pipeline results in efficient lineage reconstruction of embryos with high spatial and rotational drift. **(a)** Overview of the pipeline used for reliable automated segmentation, tracking, and lineage tracing of the imaged embryos; 1) Segmentation: low thresholds are used for the spot detection in both the green and red channel to enable detection of dimmer cells at later developmental time points. Incorrectly segmented spots are excluded by defined filters: i) exclusion of spots outside of a defined radius of the embryo, ii) replacement of incorrectly segmented double spots by one spot per one nucleus, and ii) exclusion of red spots that do not colocalize with green nuclear spots. 2) Tracking: Spatial drift as well as rapid embryo rotation complicates tracking nuclei over prolonged time windows. The segmented nuclei are used for defining reference frames based on the center of mass of the green nuclei and the orientation of the red nuclei. The alignment of the references frames of each time point compensates the spatial and rotational drifts. 3) Lineage tracing: Automated lineage tree reconstruction can make false connections when cells are dividing. By separating the calculation of the lineage trees in the photoconverted red channel from the green channel, the less complex datasets for each channel result in more consistent lineage tracing. pr-mEosFP fluorescence (green) and primed converted pr-mEosFP fluorescence (magenta) overlaid with segmentation results (green and Magenta spheres); Scale bar, 20 µm (**b**) Lineage trees from the same embryo (corresponding to Video 1 and 3) reconstructed from segmented nuclei before correction for rotational and translational drift (left), after correction for rotational and translational drift for the red channel (second left), after correction for rotational and translational drift for the green channel minus the spots that colocalize with the red spots (second right), and after final manual lineage reconstruction (right). The embryo was imaged every 15 minutes.

In a second step, the embryo was centered at its fluorescence center of mass, cropped and rotated, such that the red center of mass was oriented to the same side of the embryo in every time frame to compensate for rotational and translational drift (Figure 2a, middle column; Video 3 and 4). The resulting high-quality 5D cropped and registered datasets were on average 61±12% smaller in size (Figure 2 – figure supplement 2). The automatic tracing of a realigned embryo resulted in greatly improved lineage tracing fidelity compared to a naïve state-of-the-art lineage-tracing algorithm that was not able to reconstruct a lineage tree from rotating and spatially drifting embryos imaged with a time interval of 15 or 7.5 minutes (Bitplane Imaris cell lineage package) (Figure 2b; Figure 2 – figure supplement 3). Separating the green and red channels to generate two less complex datasets during lineage reconstruction further increased the fidelity of lineage tracing versus the dataset consisting of the green channel alone (Figure 2a, right column; Figure 2 – figure supplement 3). We assessed the power of our newly created lineage tracing algorithm by comparing the lineage trees obtained i) without corrections, ii) after embryo realignment with all algorithmic corrections, and iii) after final manual review by calculating the total distance between these lineage trees (see methods for more details)^11^. Notably, the resulting lineage trees required a minimal amount of time for manual corrections (i.e. 1-1.5 hours for the total lineage tree).

## Summary and conclusion

In summary, the presented approach enables fast, automated, high fidelity lineage tracing of mammalian pre-implantation development combined with reduced illumination time and data volume, key considerations for handling and analyzing data by the biological community^12^. In addition, the ability to correct for both spatial and rotational drift overcomes the need to exclude spinning embryos from the analysis. On a different note, it might enable the experimenter to achieve similar tracing quality with datasets acquired at lower sampling rate. In the future, implementing primed conversion to take place inside the SPIM used for volumetric imaging, will allow for repeated manual or automatic primed conversion of nuclei once the red fluorescence drops below a user-defined threshold. Similar pulse-chase experiments can then be extended even longer, ultimately being only limited by the rate of new green pr-pcFP synthesis. The combination of confined primed conversion of pr-pcFPs with our imaging pipeline will allow researchers to get more accurate insight into the dynamic processes responsible for cell fate decisions in the early mammalian embryo.

## Materials and Methods

### Molecular cloning and mRNA preparation

The coding sequences for pr-mEosFP and pCS2+-H2B-pr-EosFP were obtained by PCR amplification from pQE32-pr-mEosFP (Addgene No. 99213) and pRSET-pr-mEos2 (gift from Dominique Bourgeois) and cloned into pCS2+-H2B-Dendra2 using AgeI and SnaBI, hence replacing the Dendra2 coding sequence to obtain pCS2+-H2B-pr-EosFP (Addgene No.:XXXXX) and pCS2+-H2B-pr-Eos2. mRNA was synthesized using the mMESSAGE mMACHINE kit (ThermoFisher Scientific), followed by poly-A-tailing (ThermoFisher Scientific), and purified using a Qiagen RNAeasy kit according to manufacturer guidelines.

### mRNA microinjection of mouse preimplantation embryos and ex utero culture up to 4-cell stage

C57/Bl6 wild-type females were superovulated by hormone priming, mated to C57/Bl6 males, and mated females were euthanized by CO_2_ asphyxiation. Embryos were recovered by flushing oviducts as described previously^8,13^. Embryos were cultured at 37°C and 5% CO2 in KSOM+AA medium covered with mineral oil. mRNA constructs were microinjected into the pro-nucleus at 50 ng/ul or in both cells in two-cell stage embryos, following standard protocols. All these experiments were approved by the veterinary authority of the canton Basel Stadt, Swizerland.

### Confined primed conversion of single nuclei in mouse embryos

Confined primed conversion of single nuclei was performed on mouse embryos at the 4-cell stage as previously described in great detail^8^.

### Volumetric imaging of mouse pre-implantation embryos

Right after confined primed conversion was performed, the 4-cell stage embryos were transferred to a pre-equilibrated, inverted SPIM setup and continuously cultured/imaged until they reached blastocyst stage^2^. For each embryo a z-stack consisting of 80 planes, 3 µm apart, was acquired every 7.5 or 15 minutes.

### Mouse embryo lineage tracing

To establish a reference, mouse embryos were lineage traced using the state-of-the-art Imaris lineage tracing package (Bitplane, CH). The automated high-fidelity mouse embryo drift correction and lineage-tracing algorithm described here is explained in detail below.

### Detailed description of the automatic segmentation, registration, and lineage tracing algorithm

5D movies of photoconverted mouse embryos were processed with the following pipeline using a custom MATLAB code implemented in Imaris (Bitplane, CH).

### Cell Segmentation

1. Detect green and red cells using the Spot detector in Imaris. Use low threshold to segment all cells even at the cost of including spurious spots. Allow spot radius to be adapted to more accurately fit the volume of the segmented cell. Bright but very small spots can easily be filtered out during segmentation.

### First validation

2. Use the green spot positions to estimate the embryo diameter and discard green spots that are likely to be outside of the embryo. The radius of the embryo is roughly estimated as the median of all maximal inter-spot distances. A user-defined multiplicative factor can optionally compensate for estimation errors and prevent cells at the boundary of the embryo to be discarded should this constraint be too stringent.

3. Search for spots that occur within a small defined distance from a spot in the same channel, discard all wrongly segmented double spots on one nucleus and replace them by one new spot.

4. Discard all spurious red spots that do not colocalize with a green spot. Note that due to the equilibrium between protonated and de-protonated chromophore, green to red photoconversion of pcFPs is never exhaustive and will always retain a green population, rendering this quality control step possible.

5. A red spot discarded during the first validation can optionally be recovered if there is a valid red spot in previous time point within a user-defined search radius. This adjustment compensates for remaining miss-segmentations in the green channel.

### Embryo alignment (drift and rotational correction)/Cropping

6. Imaris Reference Frame Objects are created in MATLAB for each time point: their origin is set at the position of the center of mass (COM) of the green spots and their orientation is given by the vector 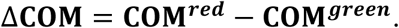 This correction still has one degree of freedom. The rotation angle around the reference frame axis is obtained by comparing the positions of the green spots at timepoints ***t*** and ***t - 1*** over 360 1-degree rotations and by choosing the angle that minimizes the cell drift between time points. The resampling is performed in Imaris.

7. Crop data to the smallest bounding volume.

### Second validation and subsetting

8. To pick up red cells that were not recovered previously, re-run the first validation on the re-aligned embryo.

9. Create a new spot object that contains the subset of green cells that do not colocalize with red cells.

### Lineage tree reconstruction

10. Imaris’ Lineage module is used to track the cells over time and reconstruct their lineage tree. The subsetting in the previous step allows us to reduce the complexity of the lineage tracing problem by breaking it down into two simpler, computationally less expansive, disjoint problems.

## Comparative Analysis of lineage trees

To assess the power of our newly created algorithm, we sought to compare the lineage trees obtained with i) no corrections, ii) after embryo realignment with all algorithmic corrections, and iii) after final manual review. We quantified the effects of the corrections and validations on the quality of the lineage trees by calculating the total distance between the lineage trees using the implementation of the tree Zhang-Shasha edit distance algorithm^11^ by Tim Henderson and Steve Johnson^14^. The zss algorithm assigns a (user-defined) cost for each node insertion, removal, and update necessary to transform an ordered tree into another, and gives therefore a quantitative measure of dissimilarity of the two trees. Small tracking differences between corrected and uncorrected trees, however, can result in quite large tree distances if the zss algorithm is applied to the complete trees. A correction that relinks one cell to its mother cell in just one time point causes the whole branch to be flagged as incorrect, and the longer the branch, the higher the distance between the trees. In other words, the earlier the tracking error occurs, the larger the distance; yet, only the first time point in the track is incorrect, and its penalty should be the same whether it happens at the beginning of the time series or the end.

To circumvent these issues, we applied the algorithm to a condensed version of the lineage trees. The condensed tree retains only the branch points of the original lineage tree (i.e. the cell divisions). Also, each branch point stores information about the original number of child nodes in its branches (i.e. the number of time points the daughter cells were tracked until their next cell division). The distance between condensed trees will flag positions where cell divisions were tracked incorrectly and tracks that have different lengths, without causing an explosion in the reported distance.

Since our acquisitions started at the 4-cell stage, we aimed to build a tree for each of the original four cells (one containing the progeny of the primed converted cell). The final, manually curated lineage was used as ground truth to quantify the effects of the various algorithmic correction steps. The sets of trees across correction schemes were assigned to each other by minimizing the spatial and temporal distance of their origins. After condensation, their pairwise distances were calculated. All distances were summed to give the total lineage tree difference. In addition, spurious trees that resulted from bad segmentation and tracking were not used for the distance calculation, since they already indirectly affected the difference of the tree from which they were erroneously detached.

## Funding

**Rubicon fellowship from the Netherlands Organisation for Scientific Research (NWO)**

- Maaike Welling

**Howard Hughes Medical Institute Janelia graduate research fellowship**

- Manuel A Mohr

**Howard Hughes Medical Institute Janelia’s Visiting Scientist Program**

- Periklis Pantazis

**Swiss National Science Foundation (SNF grant no. 31003A_144048)**

**European Union Seventh Framework Program (Marie Curie Career Integration Grant (CIG) no. 334552)**

The funding sources had no role in the study design, data collection and interpretation, or the decision to submit the work for publication.

## Acknowledgements

We thank all members of the Pantazis lab and especially M. Haffner for discussion and advice. We thank U. Nienhaus and K. Nienhaus for discussions and advice as well as E. Schreiter, and L. Looger for discussions; W.P. Dempsey for feedback on the manuscript and C. Morkunas for administrative management.

## Competing interests

P.Pa is an inventor on a patent application filed by ETH Zurich that describes primed conversion. P.Pa. and M.A.M. are inventors on a provisional patent application filed by HHMI and ETH Zurich that describes pr-mEosFP.

## Individual Author Contributions

M.A.M., and P.Pa. conceived the idea. M.W. and M.A.M planned and M.W. carried out imaging experiments with the help of M.A.M. and L.R.S. A.B. and P.L. assisted with design and execution of inverted light-sheet imaging experiments and P.Pe. performed mouse embryo injections. M.W., A.P. and M.A.M conceived and A.P. implemented the software with the help of M.W. and M.A.M. M.W. and M.A.M. wrote the manuscript with the help of P.Pa. and A.P. and editing input from the other authors. P.Pa. supervised the work.

---

**Video 1. Video of a developing embryo before drift correction**
Timelapse video of an example embryo, which shows strong spatial and rotational drift before drift correction. pr-mEosFP fluorescence (green) and primed converted pr-mEosFP fluorescence (red). Scale bars, 15 µm; framerate: one frame every 15 minutes.

**Video 2. Video of another developing embryo before drift correction.**
Timelapse video of an example embryo, which shows strong spatial and rotational drift before drift correction. pr-mEosFP fluorescence (green) and primed converted pr-mEosFP fluorescence (red). Scale bars, 10 µm; framerate: one frame every 7.5 minutes.

**Video 3. Video of a developing embryo (same as in Video 1) after drift correction.**
Timelapse video of the example embryo from Video 1 after drift correction. pr-mEosFP fluorescence (green) and primed converted pr-mEosFP fluorescence (red). Corresponding lineage trees are displayed in Figure 2d. Scale bars, 15 µm; framerate: one frame every 15 minutes.

**Video 4. Video of a developing embryo (same as in Video 2) after drift correction.** Timelapse video of the example embryo from Video 2 after drift correction. pr-mEosFP fluorescence (green) and primed converted pr-mEosFP fluorescence (red). Corresponding lineage trees are displayed in Figure 2- figure supplement 3. Scale bars, 10 µm; framerate: one frame every 7.5 minutes.

## Supplementary files include

Figure 1 – figure supplement 1: Embryos expressing H2B-pr-mEosFP develop normally.

Figure 1 – figure supplement 2: Visualizing the lineage of a single cell up to the blastocyst stage using primed converted at the 4-cell stage.

Figure 2 – figure supplement 1: Dual labeling facilitates segmentation in dense environments. Figure 2 – figure supplement 2: Embryo dataset size before and after registration.

Figure 2 – figure supplement 3: Comparison of lineage tracing results.

Supplemental methods: Detailed description of the automatic segmentation, registration, and lineage tracing algorithm

